# Machine learning to classify animal species in camera trap images: applications in ecology

**DOI:** 10.1101/346809

**Authors:** Michael A. Tabak, Mohammad S. Norouzzadeh, David W. Wolfson, Steven J. Sweeney, Kurt C. VerCauteren, Nathan P. Snow, Joseph M. Halseth, Paul A. Di Salvo, Jesse S. Lewis, Michael D. White, Ben Teton, James C. Beasley, Peter E. Schlichting, Raoul K. Boughton, Bethany Wight, Eric S. Newkirk, Jacob S. Ivan, Eric A. Odell, Ryan K. Brook, Paul M. Lukacs, Anna K. Moeller, Elizabeth G. Mandeville, Jeff Clune, Ryan S. Miller

## Abstract

1. Motion-activated cameras (“camera traps”) are increasingly used in ecological and management studies for remotely observing wildlife and have been regarded as among the most powerful tools for wildlife research. However, studies involving camera traps result in millions of images that need to be analyzed, typically by visually observing each image, in order to extract data that can be used in ecological analyses.

2. We trained machine learning models using convolutional neural networks with the ResNet-18 architecture and 3,367,383 images to automatically classify wildlife species from camera trap images obtained from five states across the United States. We tested our model on an independent subset of images not seen during training from the United States and on an out-of-sample (or “out-of-distribution” in the machine learning literature) dataset of ungulate images from Canada. We also tested the ability of our model to distinguish empty images from those with animals in another out-of-sample dataset from Tanzania, containing a faunal community that was novel to the model.

3. The trained model classified approximately 2,000 images per minute on a laptop computer with 16 gigabytes of RAM. The trained model achieved 98% accuracy at identifying species in the United States, the highest accuracy of such a model to date. Out-of-sample validation from Canada achieved 82% accuracy, and correctly identified 94% of images containing an animal in the dataset from Tanzania. We provide an R package (Machine Learning for Wildlife Image Classification; MLWIC) that allows the users to A) implement the trained model presented here and B) train their own model using classified images of wildlife from their studies.

4. The use of machine learning to rapidly and accurately classify wildlife in camera trap images can facilitate non-invasive sampling designs in ecological studies by reducing the burden of manually analyzing images. We present an R package making these methods accessible to ecologists. We discuss the implications of this technology for ecology and considerations that should be addressed in future implementations of these methods.

## Introduction

An understanding of species’ distributions is fundamental to many questions in ecology (MacArthur, 1984; Brown, 1995). Observations of wildlife can be used to model species distributions and population abundance and evaluate how these metrics relate to environmental conditions (Elith, Kearney, & Phillips, 2010; Tikhonov et al., 2017). However, developing statistically sound data for species observations is often difficult and expensive (Underwood, Chapman, & Connell, 2000) and significant effort has been devoted to correcting bias in more easily collected or opportunistic observation data (Royle & Dorazio, 2008; MacKenzie et al., 2017). Recently, technological advances have improved our ability to observe animals remotely. Sampling methods such as acoustic recordings, images from crewless aircraft (or “drones”), and motion-activated cameras that automatically photograph wildlife (i.e., “camera traps”) are commonly used (Blumstein et al., 2011; O’Connell et al., 2011; Getzin et al., 2012). These tools offer great promise for increasing efficiency of observing wildlife remotely over large geographical areas with minimal human involvement and have made considerable contributions to ecology (Rovero et al., 2013; Howe et al., 2017). However, a common limitation is these methods lead to a large accumulation of data - audio and video recordings and images - which must be first classified in order to be used in ecological studies predicting occupancy or abundance (Swanson et al., 2015; Niedballa et al., 2016). The large burden of classification, such as manually viewing and classifying images from camera traps, often constrains studies by reducing the sampling intensity (e.g., number of cameras deployed), limiting the geographical extent and duration of studies. Recently, machine learning has emerged as a potential solution for automatically classifying recordings and images.

Machine learning methods have been used to classify wildlife in camera trap images with varying levels of success and human involvement in the process. One application of a machine learning approach has been to distinguish empty and non-target animal images from those containing the target species to reduce the number of images requiring manual classification. This approach has been generally successful, allowing researchers to remove up to 76% of images containing non-target species (Swinnen et al., 2014). Development of methods to identify several wildlife species in images has been more problematic. Yu et al. (2013) used sparse coding spatial pyramid matching (Lazebnik, Schmid, & Ponce, 2006) to identify 18 species in images, achieving high accuracy (82%), but their approach necessitates each training image to be manually cropped, requiring a large time investment. Attempts to use machine learning to classify species in images without manual cropping have achieved far lower accuracies: 38% (Chen et al., 2014) and 57% (Gomez Villa, Salazar, & Vargas, 2017). However, more recently Norouzzadeh et al. (2018) used convolutional neural networks with 3.2 million classified images from camera traps to automatically classify 48 species of Serengeti wildlife in images with 95% accuracy.

Despite these advances in automatically identifying wildlife in camera trap images, the approaches remain study specific and the technology is generally inaccessible to most ecologists. Training such models typically requires extensive computer programming skills and tools for novice programmers (e.g., an R package) are limited. Making this technology available to ecologists has the potential to greatly expand ecological inquiry and non-invasive sampling designs, allowing for larger and longer-term ecological studies. In addition, automated approaches to identifying wildlife in camera trap images have important applications in detecting invasive species or sensitive species and improving their management.

We sought to develop a machine learning approach that can be applied across study sites and provide software that ecologists can use for identification of wildlife in their own camera trap images. Using over three million identified images of wildlife from camera traps from five locations across the United States, we trained and tested deep learning models that automatically classify wildlife. We provide an R package (Machine Learning for Wildlife Image Classification; MLWIC) that allows researchers to classify camera trap images from North America or train their own machine learning models to classify images. We also address some basic issues in the potential use of machine learning for classifying wildlife in camera trap images in ecology. Because our approach nearly eliminates the need for manual curation of camera trap images we also discuss how this new technology can be applied to improve ecological studies in the future.

## Materials and Methods

### Camera trap images

Species in camera trap images from five locations across the United States (California, Colorado, Florida, South Carolina, and Texas) were identified manually by researchers (see Appendix S1 for a description of each field location). Images were either classified by a single wildlife expert or evaluated independently by two researchers; any conflicts were decided by a third observer (Appendix S1). If any part of an animal (e.g., leg or ear) was identified as being present in an image, this was included as an image of the species. This resulted in a total of 3,741,656 classified images that included 28 species or groups (see Table 1) across the study locations. We present these images and their classifications for other scientists to use for model development as the North American Camera Trap Images (NACTI) dataset. Images were re-sized to a resolution of 256 × 256 pixels using a custom Python script before running models to increase processing speed. A subset of images (approximately 10%) was withheld using conditional sampling to be used for testing of the model (described below). This resulted in 3,367,383 images used to train the model and 374,273 images used for testing.

**Table 1:**
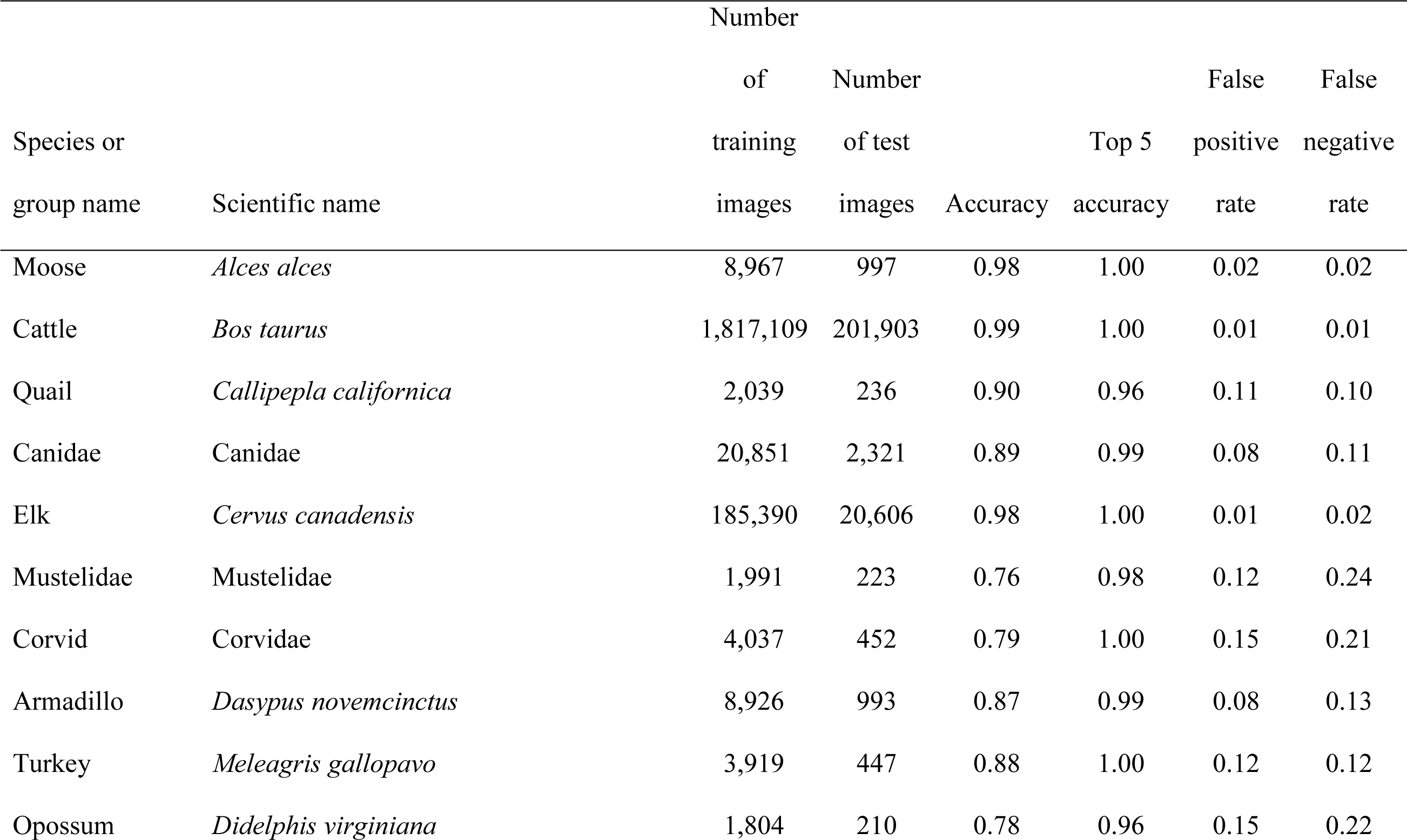
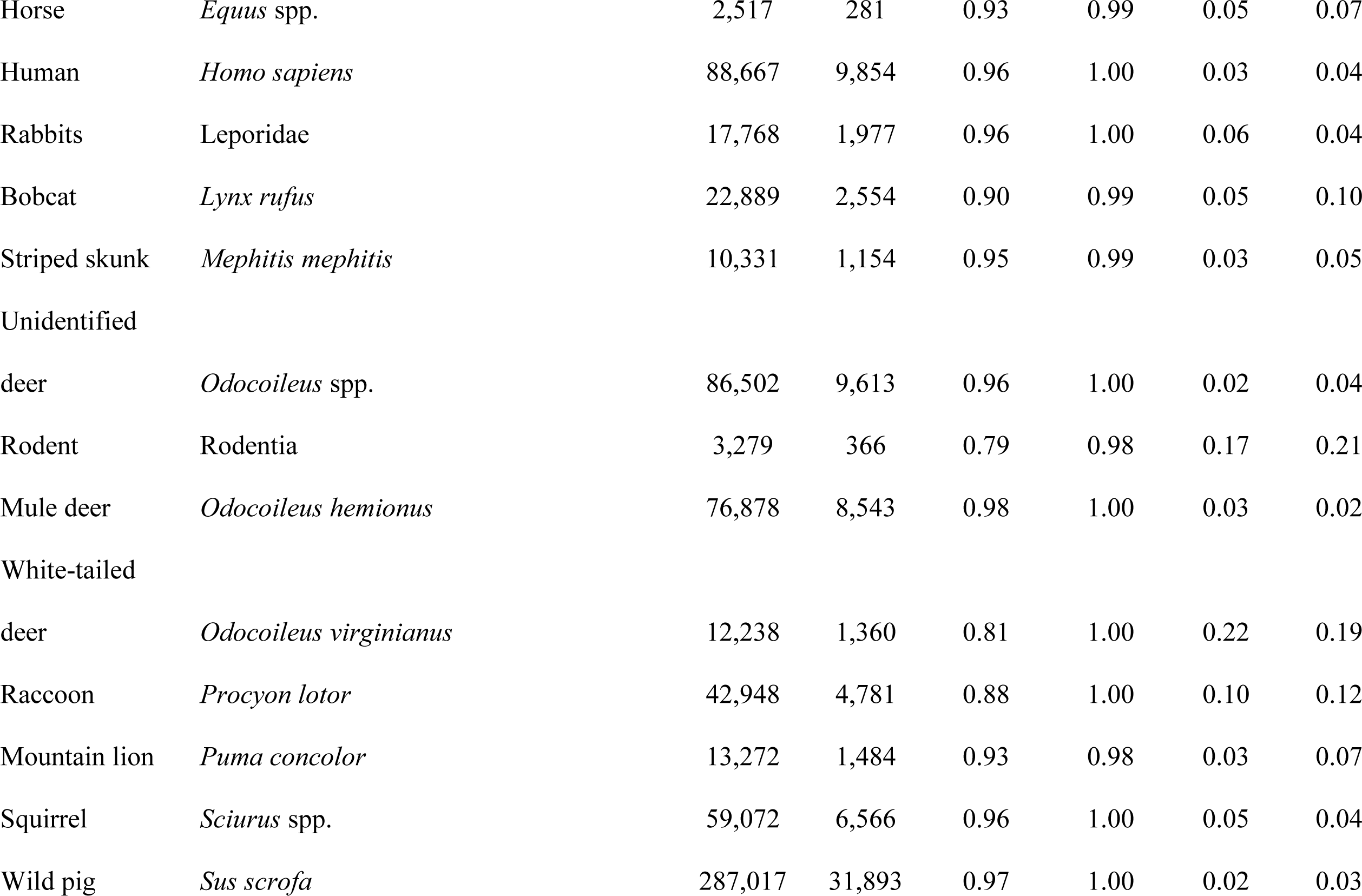
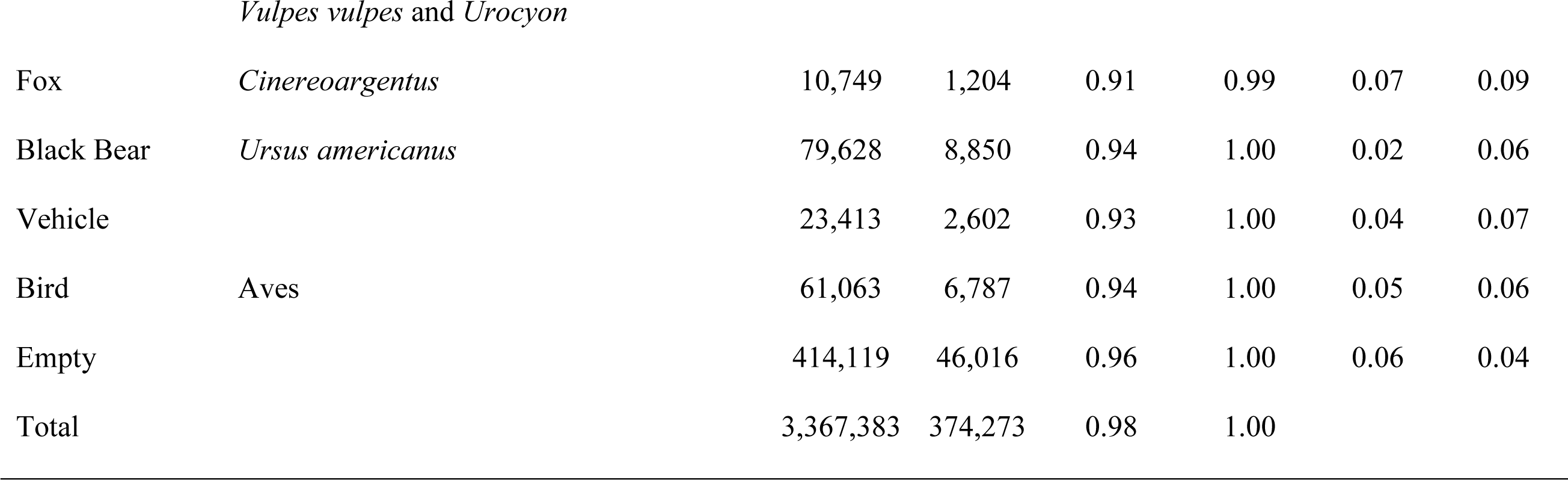
Accuracy of the Species Level model

### Machine learning process

Supervised machine learning algorithms use training examples to “learn” how to complete a task (Mohri, Rostamizadeh, & Talwalkar, 2012; Goodfellow, Bengio, & Courville, 2016). One popular class of machine learning algorithms is artificial neural network, which loosely mimics the learning behavior of the mammalian brain (Gurney, 2014; Goodfellow et al., 2016). An artificial neuron in a neural network has several inputs, each with an associated weight. For each artificial neuron, the inputs are multiplied by the weights, summed, and then evaluated by a non-linear function, which is called the activation function (e.g., Sigmoid, Tanh, or Sine). Usually each neuron also has an extra connection with a constant input value of 1 and its associated weight, called a “bias,” for neurons. The result of the activation function can be passed as input into other artificial neurons or serve as network outputs. For example, consider an artificial neuron with three inputs (*I*_1_, *I*_2_, and *I*_3_); the output (θ) is calculated based on:

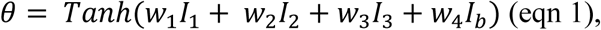

where *w*_1_, *w*_2_, *w*_3_ and *w*_4_ are the weights associated with each input, *I*_*b*_ is the bias, and *Tanh(x)* is the activation function (Fig. 1). To solve complex problems multiple neurons are needed, so we put them into a network. We arrange neurons in a hierarchical structure of layers; neurons in each layer take input from the previous layer, process them, and pass the output to the next layer. Then, an algorithm, called backpropagation (Rumelhart, Hinton, & Williams, 1986), tunes the parameters of the neural network (weights and bias values) enabling it to produce the desired output when we feed an input to the network. This process is called training. To adjust the weights, we define a loss function as a measure of the difference between the predicted (current) output of the neural network and the correct output *(Y).* The loss function *(L)* is the mean squared error:

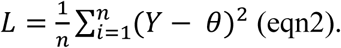

We compute the contribution of each weight to the loss value 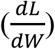 using the chain rule in calculus. Weights are then adjusted so the loss value is minimized. In this “weight update” step, all the weights are updated to minimize *L:*

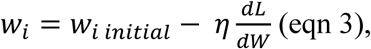

where *η* is the learning rate and is chosen by the scientist. A higher *η* indicates larger steps are taken per training sample, which may be faster, but a value that is too large will be imprecise and can destabilize learning. After adjusting the weights, the same input should result in an output that is closer to the desired output. For more details of backpropagation and training, see Goodfellow et al., 2016.

**Figure 1:**
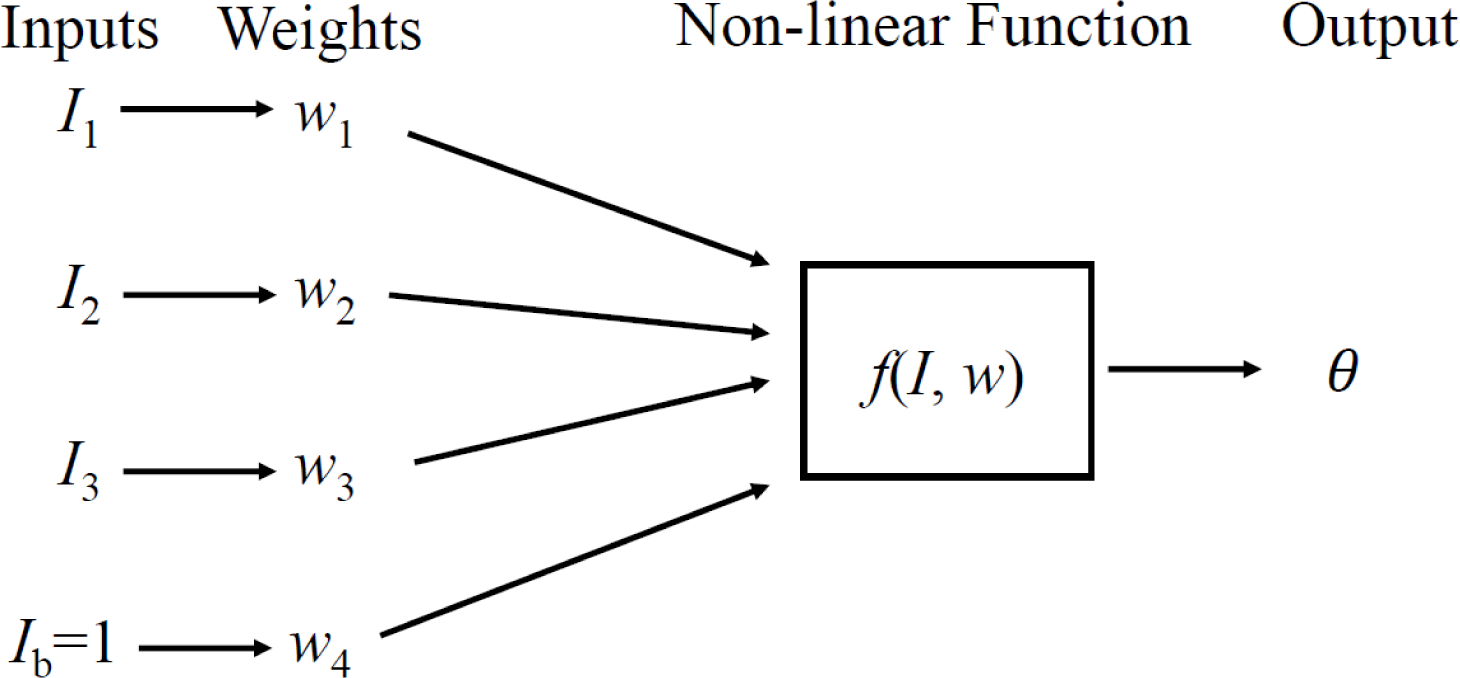
Within an artificial neural network, inputs (*I*) are multiplied by their weights (*w*), summed, and then evaluated by a non-linear function, which also accounts for bias *(l*_*b*_). The output *(θ)* can be passed as input into other neurons or serve as network outputs. Backpropagation involves adjusting the weights so that a model can provide the desired output.

In fully connected neural networks, each neuron in every layer is connected to (provides input to) every neuron in the next layer. Conversely, in convolutional neural networks, which are inspired by the retina of the human eye, several convolutional layers exist in which each neuron only receives input from a small sliding subset of neurons (“receptive field”) in the previous layer. We call the output of a group of neurons the “feature map,” which depicts the response of a neuron to its input. When we use convolutional neural networks to classify animal images, the receptive field of neurons in the first layer of the network is a sliding subset of the image. In subsequent layers, the receptive field of neurons is a sliding subset of the feature map from previous layers. We interpret the output of the final layer as the probability of the presence of species in the image. A softmax function is used at the final layer to ensure that the outputs sum to one. For more details on this process, see Simonyan & Zisserman, 2014.

Deep neural networks (or “deep learning”) are artificial networks with several (> 3) layers of structure. In our example, we provided a set of animal images from camera traps of different species and their labels (species identifiers) to a deep neural network, and the model learned how to identify species in other images that were not used for training. Once a model is trained, we can use it to classify new images. The trained model uses the output of the final layer in the network to assign a confidence to each species or group it evaluates, where the total confidence assigned to all groups for each image sums to one. Generally, the majority of the confidence is attributed to one group, the “top guess.” For example, for 90% of the images in our test dataset, the model attributed > 95% confidence to the top guess. Therefore, for the purpose of this paper, we mainly discuss accuracy with regard to the top guess, but our R package presents the five groups with the highest confidence, the “top five guesses,” and the confidence associated with each guess.

Neural network architecture refers to several details about the network including the type and number of neurons and the number of layers. We trained a deep convolutional neural network (ResNet-18) architecture because it has few parameters, but performs well; see He et al. (2016) for full details of this architecture. Networks were trained in the TensorFlow framework (Adabi et al., 2016) using Mount Moran, a high performance computing cluster (Advanced Research Computing Center, 2012). First, since invasive wild pigs *(Sus scrofa)* are a subject of several of our field studies, we developed a “Pig/no pig” model, in which we determined if a pig was either present or absent in the image. In the “Species Level” model, we identified images to the species level when possible. Specifically, if our classified image dataset included < 2,000 images for a species, it was either grouped with taxonomically similar species (by genera, families, or order), or it was not included in the trained model (Table 1). In the “Group Level” model, species were grouped with taxonomically similar species into classifications that had ecological relevance (Appendix S2). The Group Level model contained fewer groups than the Species Level model, so that more training images were available for each group. We used both models because if the Species Level model had poor accuracy, we predicted the Group Level model would have better accuracy since more training images would be available for several groups. As it is the most broadly applicable model and is the one implemented in the MLWIC package, we will mainly discuss the Species Level model here, but show results from the Group Level to demonstrate alternative approaches.

For each of the three models, 90% of the classified images for each species or group were used to train the model and 10% of the images were used to test it in most cases. However, we wanted to evaluate the model’s performance for each species present at each study site, so we altered training-testing allocation for the rare situations where there were few classified images of a species at a site. Specifically, with 1-9 classified images for a species at a site, we used all of these images for testing and none for training; for site-species pairs with 10-30 images, 50% were used for training and testing; and for > 30 images per site for each species, 90% were allocated to training and 10% to testing (Appendices S3 - S7 show the number of training and test images for each species at each site).

### Evaluating model accuracy

Model testing was conducted by running the trained model on the withheld images that were not used to train the model. Accuracy (*A*) was assessed as the proportion of images in the test dataset (*N*) that were correctly classified (*C*) by the top guess (*A = C/N*). Top 5 accuracy (*A*5) was defined as the proportion of images in the test dataset that were correctly classified by any of the top 5 assignments (*C*5; *A*5 = *C*5/*N*). For each species or group we calculated the rate of false positives (*FP*) as the proportion of images classified as this species or group (*N*_*modei group*_) by the model’s top guess that contained a different species according to human observers (*N*_*true other*_; *FP* = *N*_*true other*_/*N*_*model group*_). We calculated the rate of false negatives for each species (*FN*) as the proportion of images observers classified as a specific species or group (*N*_*true group*_) that the model’s top guess classified differently (*N*_*modei other*_; *FN* = *N*_*model other*_/*N*_*true group*_). This assumes the observers were correct in their classification of images. We fit generalized additive models (GAMs) to the relationship between accuracy and the logarithm (base 10) of the number of images used to train the model. We also calculated the accuracy and rates of error specific to each of the five data sets from which images were acquired.

To evaluate how the model would perform for a completely new study site in North America, we used a dataset of 5,900 classified images of ungulates (moose, cattle, elk, and wild pigs) from Saskatchewan, Canada by running the Species Level model on these images. We also evaluated the ability of the model to operate on images with a completely different species community (from Tanzania) to determine the model’s ability to correctly classify images as having an animal or being empty when encountering new species that it has not been trained to recognize. This was done using 3.2 million classified images from the Snapshot Serengeti dataset (Swanson et al., 2015).

## Results

Our models performed well, achieving ≥ 97.5% accuracy of identifying the correct species with the top guess (Table 2). The model determining presence or absence of wild pigs had the highest accuracy of all of our models (98.6%; Pig/no pig; Table 2). For the Species Level and Group Level models, the top 5 accuracy was > 99.9%. The model confidence in the correct answer varied, but was mostly > 95%; see Fig. 2 for confidences for each image for three example species. Supporting a similar finding for camera trap images in Norouzzadeh et al. (2018), and a general trend in deep learning (Goodfellow et al., 2016), species and groups that had more images available for training were classified more accurately (Fig. 3, Table 1). GAMs relating the number of training images with accuracy predicted 95% accuracy could be achieved when approximately 71,000 training images were available for a species or group. However, these models were not perfect fits to the data, and for several species and groups, 95% accuracy was achieved with fewer than 70,000 images (Fig. 3). We found there was not a large effect of daytime vs. nighttime on accuracy in the Species Level model as daytime accuracy was 98.2% and nighttime accuracy was 96.6%. The top 5 accuracies for both times of day were ≥ 99.9%. When we subsetted the testing dataset by study site, we found that site-specific accuracies ranged from 90-99% (Appendices S3 - S7). The model performed poorly (0 - 22% accuracy) for species in the four instances when the model did not include training images from that site (when < 10 classified images were available for the species/study site combination; Appendices S3 - S7).

**Table 2:** Accuracy (across all images for all species) of the three deep learning tasks analyzed

| Model | Accuracy (%) |
| --- | --- |
| Pig/no pig | 98.6 |
| Species Level | 97.5 |
| Group Level | 97.8 |

**Fig. 2:**
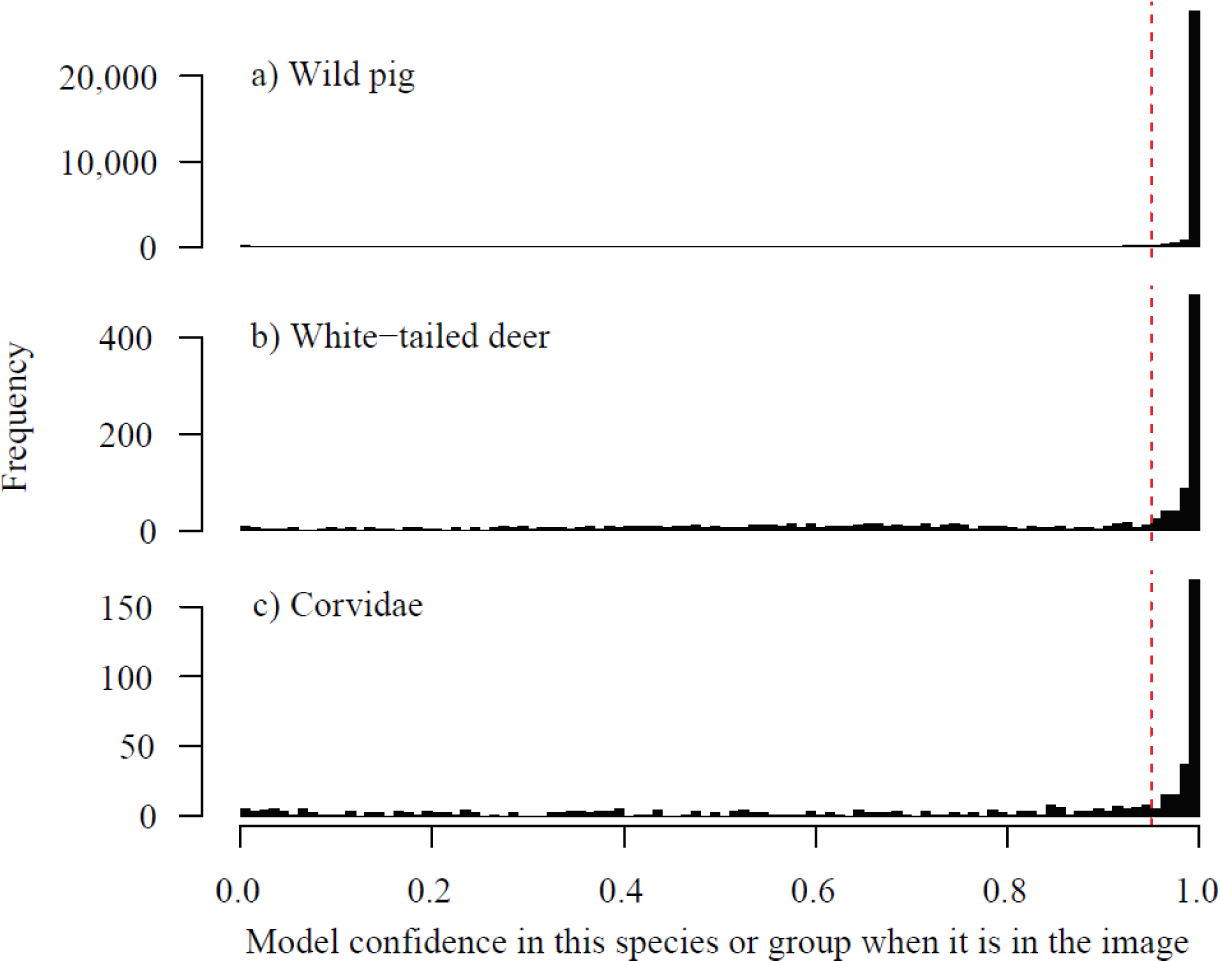
Histograms represent the confidence assigned by all of the top five guesses by the Species Level model for each of these three example species when it was present in an image. The dashed line represents 95% confidence; the majority of model-assigned confidences were greater than this value.

**Fig. 3:**
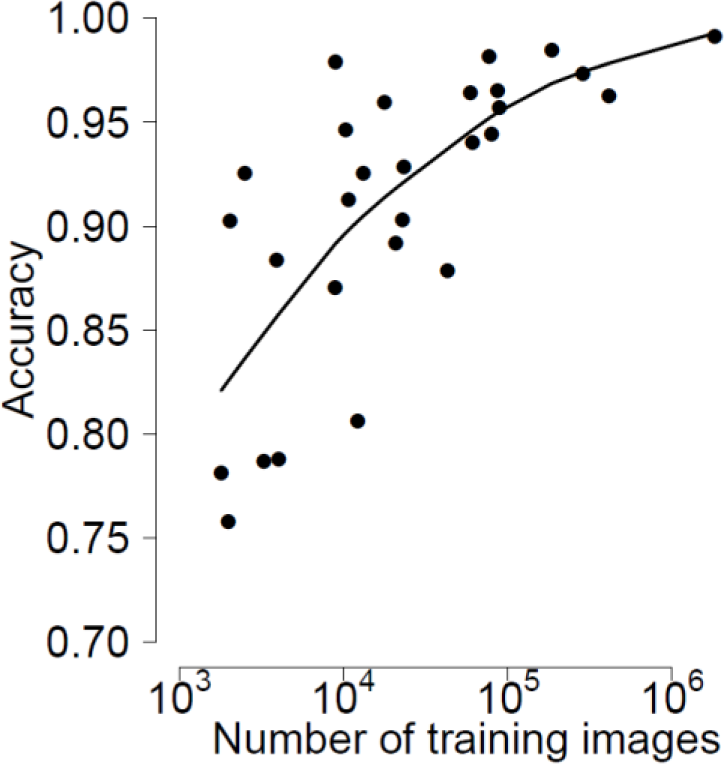
Machine learning model accuracy increased with the size of the training dataset. Points represent each species or group of species. The line represents the result of generalized additive models relating the two variables.

Upon further investigation, we found these images were difficult to classify manually. For example, striped skunks in Florida were misclassified in both of the images from this study site (Appendix S5). These images both contained the same individual at the same camera, and most wildlife experts would not classify it as a skunk (Appendix S8).

When we conducted out-of-sample validation by using our model to evaluate images of ungulates from Canada, we achieved an overall accuracy of 81.8% with a top 5 accuracy of 90.9%. When we tested the ability of our model to accurately predict presence or absence of an animal in the image using the Serengeti Snapshot dataset, we found that 85.1% were classified correctly as empty, while 94.3% of images containing an animal were classified as containing an animal. Our trained model was capable of classifying approximately 2,000 images per minute on a Macintosh laptop with 16 gigabytes (GB) of RAM.

## Discussion

To our knowledge, our Species Level model achieved the highest accuracy (97.5%) to date in using machine learning for wildlife image classification (a recent paper achieved 95% accuracy; Norouzzadeh et al., 2018). This model performed almost as well during the night as during the day (accuracy = 97% and 98%, respectively). We provide this model as an R package (MLWIC), which is especially useful for researchers studying the species and groups available in this package (Table 1) in North America, as it performed well in classifying ungulates in an out-of-sample test of images from Canada. The model can also be valuable for researchers studying other species by removing images without any animals from the dataset before beginning manual classification, as we achieved high accuracy in separating empty images from those containing animals in a dataset from Tanzania. This R package can also be a valuable tool for any researchers that have classified images, as they can use the package to train their own model that can then classify any subsequent images collected.

### Optimizing camera trap use and application in ecology

The ability to rapidly identify millions of images from camera traps can fundamentally change the way ecologists design and implement wildlife studies. Camera trap projects amass large numbers of images which require a sizable time investment to manually classify. For example, the Snapshot Serengeti project (Swanson et al., 2015) amassed millions of images and employed 28,000 volunteers to manually classify 1.5 million images (Swanson et al., 2016; Palmer et al., 2017). We found researchers can classify approximately 200 images per hour. Therefore, a project that amasses 1 million images would require 10,000 hours for each image to be doubly observed. To reduce the number of images that need to be classified manually, ecologists using camera traps often limit the number of photos taken by reducing the size of camera arrays, reducing the duration of camera trap studies, and imposing limits on the number of photos a camera takes (Kelly et al., 2008; Scott et al., 2018). This constraint can be problematic in many studies, particularly those addressing rare or elusive species that are often the subject of ecological studies (O’Connell et al., 2011), as these species often require more effort to detect (Tobler et al., 2008). Using deep learning methods to automatically classify images essentially eliminates one of the primary reasons camera trap arrays are limited in size or duration. The Species Level model in our R package can accurately classify 1 million images in less than nine hours with minimal human involvement.

Another reason to limit the number of photos taken by camera traps is storage limitations on cameras (Rasambainarivo et al., 2017; Hanya et al., 2018). When classifying images manually, we might try to use high resolution photos to improve technicians’ abilities to accurately classify images, but higher resolution photos require more storage on cameras. Our results show a model can be accurately trained and applied using low-resolution (256 × 256 pixel) images, but many of these images were re-sized from a higher resolution, which might contain more information than those which originated at a low resolution. Nevertheless, we hypothesize a model can be accurately trained using images from low resolution cameras, and our R package allows users who have such images to test this hypothesis. If supported, this can make camera trap data storage much more efficient. Typical cameras set for 2048 × 1536 pixel resolution will run out of storage space when they reach approximately 1,250 photos per GB of storage. Taking low resolution images instead can increase the number of photos stored per GB to about 10,000 and thus decrease the frequency at which researchers must visit cameras to change storage cards by a factor of eight. Minimizing human visitation also will reduce human scents and disturbances that could deter some species from visiting cameras. In the future, it may be possible to implement a machine learning model on a game camera (Elias et al., 2017) that automatically classifies images as empty or containing animals so that empty images are discarded immediately and not stored on the camera. This type of approach could dramatically reduce the frequency with which technicians need to visit cameras. Furthermore, if models effectively use low-resolution images, it is not necessary for researchers to purchase high resolution cameras. Instead, researchers can purchase lower cost, lower resolution cameras and allocate funding toward purchasing more cameras and creating larger camera arrays.

### Applications to management of invasive and sensitive species

By removing some of the major burdens associated with the use of camera traps, our approach can be utilized by ecologists and wildlife managers to conduct more extensive camera trapping surveys than were previously possible. One potential use is in monitoring the distribution of sensitive or invasive species. For example, the distribution of invasive wild pigs in North America is commonly monitored using camera traps. Humans introduce this species into new locations that are often geographically distant from their existing range (Tabak et al., 2017), which can quickly lead to newly-established populations. Camera traps could be placed in areas at risk for introduction and provide constant surveillance. An automated image classification model that simply ‘looks’ for pigs in images could monitor camera trap images and alert managers when images with pigs are found, facilitating removal of animals before populations establish. Additionally, after wild pigs have been eradicated from a region, camera traps could be used to monitor the area to verify eradication success and automatically detect re-colonization or reintroduction events. Similar approaches can be used in other study systems to more rapidly detect novel invasive species arrivals, track the effects of management interventions, monitor species of conservation concern, or monitor sensitive species following reintroduction efforts.

### Limitations

Using out-of-sample model validation on a dataset from Canada revealed a lower accuracy (82%) than at study sites from which our model was trained. Additionally, when we did not include images of species/site combinations in training the model, due to low sample sizes, the model performed poorly (Appendices S3 - S7; but these images were often difficult to classify even by wildlife experts, Appendix S8). One potential explanation is the model evaluated both the animal and the environment in the image and these are confounded in the species identification (Norouzzadeh et al., 2018). Therefore, the model may have lower accuracies in environments that were not in the training dataset. Ideally, the training dataset would include training images representing the range of environments in which a species exists. Our model includes training images from diverse ecosystems, making it relevant for classifying images from many locations in North America. A further limitation is in our reported overall accuracy, which is reported across all of the images that were available for testing, and we had considerable imbalance in the number of images per species (Table 1). We provide accuracies for each species, so the reader can more directly inspect model accuracy. Finally, our model was trained using images that were classified by human observers, which are capable of making errors (O’Connell et al., 2011; Meek, Vernes, & Falzon, 2013), meaning some of the images in our training dataset were likely misclassified. Supervised machine learning algorithms require such training examples, and therefore we are unaware of a method for training such models without the potential for human classification error. Instead, we must acknowledge that these models will make mistakes due to imperfections in both human observation and model accuracy.

### Future directions

As this new technology becomes more widely available, ecologists will need to decide how it will be applied in ecological analyses. For example, when using machine learning model output to design occupancy and abundance models, we can incorporate accuracy estimates that were generated when conducting model testing. The error of a machine learning model in identifying a species is similar to the problem of imperfect detection of wildlife when conducting field surveys. Wildlife are often not detected when they are present (false negatives) and occasionally detected when they are absent (false positives); ecologists have developed models to effectively estimate occupancy when data have these types of errors (Royle & Link, 2006; Guillera-Arroita et al., 2017). We can use Bayesian occupancy and abundance models where the central tendencies of the prior distributions for the false negative and false positive error rates are derived from testing the machine learning model (e.g., values in Table 1). While we would expect false positive rates in occupancy models to resemble the false positive error rates for the machine learning model, false negative error rates would be a function of the both the machine learning model and the propensity for some species to avoid detection by cameras when they are present (Tobler et al., 2015).

Another area in need of development is how to group taxa when few images are available for the species. We grouped species when few images were available for model training using an arbitrary cut off of approximately 2,000 images per group (Table 1). We had few images of horses *(Equus* spp.), but the model identified these images relatively well (93% accuracy), presumably because they are phenotypically different from other species in our dataset. We also had few images of opossums *(Didelphis virginiana)*, but we did not group this species because it is phenotypically different from other species in our dataset and was of ecological interest in our studies; we achieved lower accuracy for this species (78%). We also included a group for rodents from species for which we only had few images *(Erethizon dorsatum, Marmota flaviventris, Genomys* spp., *Mus* spp., *Neotoma* spp., *Peromyscus* spp., *Tamais* spp., and *Rattus* spp.). The model achieved relatively low accuracy for this group (79%), presumably because there were few images for training (3,279) and members of this group are phenotypically different, making it difficult for the model to train on this group. When researchers develop new machine learning models, they will need to consider the available data, the species or groups in their study, and the ecological question that the model will help address.

Here, we mainly focused on the species or class that the model predicted with the highest confidence (the top guess), but in many cases researchers may want to incorporate information from the model’s confidence in the guess and additional model guesses. For example, if we are interested in the highest overall accuracy, we could only consider images where the confidence in the top guess is > 95%. If we subset the results from our model test in this manner, we remove 10% of the images, but total accuracy increases to 99.6%. However, if the objective of a project is to identify rare species, researchers may want to focus on all images in which the model predicts that species to be in the top 5 guesses (the 5 species or groups that the model predicts to have the highest confidence). In our model test, the correct species was in the top 5 guesses in 99.9% of the images, indicating that this strategy may be viable.

We expect the performance of machine learning models to improve in the future (Jordan & Mitchell, 2015), allowing ecologists to further exploit this technology. Our model required manual identification of many images to obtain high levels of accuracy (Table 1). Our model was also limited in that we were only able to classify the presence or absence of species; we were not able to determine the number of individuals, their behavior, or demographics. Similar machine learning models are capable of including the number of animals and their behavior in classifications (Norouzzadeh et al., 2018), but we could not include these factors because they were rarely recorded manually in our dataset. As machine learning techniques improve, we expect models will require fewer manually classified images to achieve high accuracy in identifying species, counting individuals, and specifying demographic information. Furthermore, as scientists begin projects intending to use machine learning to classify images, they may be more willing to spend time extracting detailed information from fewer images instead of obtaining less information from all images. This development would create a larger dataset of information from images that can be used to train models. As machine learning algorithms improve and ecologists begin considering this technology when they design studies, we think that many novel applications will arise.

As camera trap use is a common approach to studying wildlife worldwide, there are likely now large datasets of classified images. If scientists work together and share these datasets, we can create large image libraries that span continents (Steenweg et al., 2017); we may eventually be able to train a machine learning model that can identify many global species and be used by researchers globally. Further, effectively sharing images and classifications can potentially be integrated with a web-based platform, similar to that employed by Camera Base (http://www.atrium-biodiversity.org/tools/camerabase) or eMammal (https://emammal.si.edu/).

## Acknowledgements

We thank the hundreds of volunteers and employees who manually classified images and deployed camera traps. We thank Dan Walsh for facilitating cooperation amongst groups. Camera trap projects were funded by the U.S. Department of Energy under award # DE-EM0004391 to the University of Georgia Research Foundation; USDA Animal and Plant Health Inspection Service, National Wildlife Research Center and Center for Epidemiology and Animal Health; Colorado Parks and Wildlife; Canadian Natural Science and Engineering Research Council; University of Saskatchewan; and Idaho Department of Game and Fish.

## Data Accessibility

The trained Species Level model is available in the R package MLWIC from github (https://github.com/mikeyEcology/MLWIC). We provide the 3.7 million classified images as the North American Camera Trap Images (NACTI) dataset in a digital repository.

## Author Contributions

MAT, RSM, KCV, NPS, SJS, and DWW conceived of the project; DWW, JSL, MAT, RKB, BW, PAD, JCB, MDW, BT, PES, NPS, KCV, JMH, ESN, JSI, EAO, RKB, PML, and AKM oversaw collection and manual classification of wildlife in camera trap images from the study sites; MSN and JC developed and programmed the machine learning models; MAT led the analyses and writing of the R package; EGM assisted with R package development and computing; MAT and RSM led the writing. All authors contributed critically to drafts and gave final approval for submission.

## Supporting Information

**Appendix S1.** Site descriptions for each of the study locations

**Appendix S2.** Accuracy of the Group Level for each species

**Appendix S3.** Accuracy of the Species Level model at the Tejon research site in California.

**Appendix S4.** Accuracy of the Species Level model in Colorado

**Appendix S5.** Accuracy of the Species Level model at Buck Island Ranch in Florida

**Appendix S6.** Accuracy of the Species Level model at the Camp Bullis Military Training Center in Texas

**Appendix S7.** Accuracy of the Species Level model at the Savannah River Ecology Laboratory in South Carolina

**Appendix S8.** Image classified as a striped skunk by humans, but cattle by the Species Level model

